# Assessing the potential of germplasm collections for the management of genetic diversity: the case of the French National Cryobank

**DOI:** 10.1101/2023.07.19.549644

**Authors:** Alicia Jacques, Delphine Duclos, Coralie Danchin-Burge, Marie-José Mercat, Michèle Tixier-Boichard, Gwendal Restoux

## Abstract

Through a combination of selective pressure and genetic drift, there has been a notable erosion of genetic diversity in domesticated animal populations. In response, many countries, including France, have developed gene banks in order to conserve reproductive genetic material. Cryopreserved resources can potentially be used to manage the genetic diversity of populations, but this opportunity is seldom exploited. As an initial step toward this goal, we describe here a methodology for the characterization of the genetic diversity of cryopreserved collections. Using the example of the French National Cryobank, this study employs newly proposed biodiversity metrics to conduct a detailed assessment of the status of collections for six livestock species: cattle, sheep, goat, horse, donkey, and pig.

Both the amount of resources available and their characteristics varied among species and/or breeds. Overall, breeds with a large commercial distribution had more donors in the collection than local breeds, while the number of doses available was mainly determined by the physiology of the species. An adapted version of the Gini-Simpson equitability index revealed an unbalanced number of donors between breeds for some species. Similarly, estimates of effective donor numbers (De) highlighted the unequal distribution of donors within a breed. Finally, we developed a new index of diversity impact (IDI) to assess the potential of a collection to reintroduce genetic diversity in contemporary populations. The IDI was calculated on the basis of pedigree data for 17 breeds of three livestock species, pig, sheep, and cattle, which differed in both population size and management program. IDI values are negative when the use of cryoconserved sires would decrease the overall kinship of the current population and positive when it would increase it, enabling the most interesting donors to be chosen for immediate use. Negative (favorable) IDI values 41 were found for both local breeds as well as for commercial populations. In general, older collections exhibited better IDI values but recently collected donors could also be useful for populations undergoing strong selection. Within a breed, IDI can be computed individually and thus be used to select the best sires for a given objective. In the absence of pedigree data, IDI values could also be calculated on the basis of marker genotypes.

Overall, this study proposes a framework for the assessment of germplasm collections in the service of various objectives. Compared to FAO indicators motivated by breed reconstitution, the Gini-Simpson and De indices can help to plan sampling more efficiently, whereas IDI can guide donor selection in order to manage the diversity of existing populations. These indicators can be calculated at regular intervals to support the planning and management of collections at national and international levels and help population managers to exploit the resources currently available.

## Introduction

In recent years, a drastic decrease in biodiversity has been observed in animal populations. This is especially true for domestic animals, in which a large number of breeds worldwide have become endangered (FAO, 2015) and the amount of within-breed genetic diversity has declined in numerous species.

In light of the major environmental changes currently in progress (Hoffmann, 2010) and the significant challenges posed by the agro-ecological transition (Dumont et al., 2013; Kantanen et al., 2015; Ducos et al., 2021), it is critical that livestock populations maintain their genetic diversity in order to ensure their capacity to adapt to future environmental changes. In closed populations of domestic animals, a reduction in genetic diversity is inevitable, but it is important for population managers to limit this loss over time. Rare breeds—including local breeds with limited population sizes (Scherf, 2000; Leroy et al., 2013)— are further threatened by unpredictable changes in allele frequencies caused by genetic drift, which is stronger when the effective population size is smaller (Falconer & Mackay, 1996; Willi et al., 2006). Moreover, in the case of breeds under selection, years of artificial selection have led to an increase in inbreeding, especially for populations exhibiting an unbalanced use of sires. Although selection aims to improve performance with regard to a defined objective, any potential response to selection is constrained by the amount of genetic variation available in the population. Measures must therefore be taken to limit this decline and to promote the genetic diversity of livestock populations.

In order to preserve the genetic diversity of farm animals, many countries have set up gene banks to conserve their germplasms (Mara et al., 2013; Blackburn, 2018). Over the years, these organizations have created valuable collections for *ex situ* conservation of animal genetic resources, offering opportunities to support the conservation of endangered livestock breeds and to manage the genetic diversity of domestic animal populations *in situ*. Frozen materials such as embryos, oocytes, or semen can restore allelic variants that have been lost over time due to selection and genetic drift. For instance, the work of Eynard *et al*. (2018), Doekes *et al*. (2018b), and Jacques *et al*. (2023) all demonstrated that the use of cryopreserved genetic resources could reintroduce genetic diversity to breeding programs, especially when there are concomitant changes in the breeding goals (Leroy *et al*., 2011).

In order to evaluate the potential of cryopreserved genetic resources, it is first necessary to identify the main features of the stocks in the cryobank. For example, 50% of the collections in the US National Gene Bank represent rare breeds with small population sizes and do not necessarily meet the requirements to reconstitute a breed, whereas collections for breeds with larger populations tend to be more complete (Blackburn et al., 2019). For Dutch cattle, collections have been partially optimized to maximize diversity within seven native breeds, and their gene bank could therefore play a major role in the conservation and maintenance of these breeds over time (van Breukelen et al., 2019). A comparison between the Dutch, French, and US germplasm collections for Holstein-Friesian cattle revealed genetic similarities in part of the collections but also notable divergences, with evidence of the breed’s ancient origins only present in the US collections (Danchin-Burge et al., 2011). Finally, for eight French dairy cattle breeds that were exhibiting a decline in genetic diversity, an investigation of the collections in the French National Cryobank (FNC) revealed a lower relatedness to current females for cryopreserved bulls than for contemporary males, demonstrating the effectiveness of the FNC sampling procedure (Danchin-Burge et al., 2012).

France contains a large diversity of breeds from different animal species, and these breeds differ greatly in terms of both population size and management program. For example, Holstein and Montbéliarde are the two leading French dairy breeds with, respectively, nearly 3,400,000 and 800,000 females born between 2016 and 2019. In comparison, the local breeds Abondance and Tarentaise had about 50,000 and 17,000 females born over the same period, respectively (Danchin-Burge, 2020). As of 2015, the French agriculture ministry had officially recognized 158 breeds from six domestic mammal species, but the majority of the 115 local French breeds are considered threatened (Verrier et al., 2015). Although conservation programs exist, their implementation and sustainability depend on many factors, including socio-cultural considerations (Lauvie et al., 2011). Altogether, the local French breeds represent a wealth of biodiversity and often have particular characteristics or adaptations that add value to certain territories and are important to preserve (INRAE, 2023). In this context, the French National Cryobank was created in 1999 with the goal of preserving the reproductive material of different breeds of domestic animals. Since 2012, these efforts have been complemented by the work of the CRB-Anim project (French Biological Animal Resource Center, https://www.crb-anim.fr), which aims to fill the gaps and enrich the collections of the FNC (CRB-Anim, https://doi.org/10.15454/1.5613785622827378E12). Currently, there is material from 21 species and more than 8000 donors stored in the FNC, representing a valuable resource for the management of current populations.

However, the use of cryopreserved materials from the FNC (Additional file 1: Table S1 and Additional file 2: Figure S1) remains quite limited, and incentives to encourage their use are therefore to be proposed. A particular problem is that, beyond FAO recommendations on the reconstitution of an extinct breed (FAO, 2012), there is currently no framework for assessing or evaluating the potential of germplasm collections. Consequently, the present study aimed to i) define a framework for the evaluation and comparison of the diversity contained within collections, and ii) apply this framework to characterize the current collections of the FNC and assess their potential for breed diversity management.

## Methods

### General descriptors

Data on donor animals were obtained from the FNC database. We focused on six mammalian species of farm animals that represent the majority of FNC collections: cattle, sheep, goat, pig, horse, and donkey. For each donor, we retrieved its national identifier, breed, year of birth, the amount of material (either straws or pellets) available in the collection, the main motivation for entry into collection, and the year the donor was first collected, if available.

The motivation for entry into the collection was defined at the time of the creation of the FNC as one of three main categories (types): type I for endangered breeds, type II for individuals with one or more exceptional characteristics, and type III for individuals who were representative of their breed at a particular time.

For all donors, we calculated the number of doses available, with a dose being defined as the species-specific number of frozen semen straws used for one artificial insemination. For ruminants one dose corresponds to one straw, while for horses and donkeys one dose corresponds to eight straws. For pigs, the dose differs according to the freezing protocol; one dose may be composed of five or ten straws (depending on semen processing facilities), or of a few dozen pellets for semen frozen before 1996.

We also characterized the different breeds based on their diffusion (Diff) and use (Use). We defined four values for diffusion: ‘Local’ for breeds present in a single area, ‘National’ for breeds raised all over France, ‘International’ for transboundary breeds in which animals or their reproductive materials are traded internationally, or ‘Not Available’. The use factor had six potential values depending on the main use of the breed: Meat, Dairy (milk or cheese), Wool, Riding_horse, Draft_horse, or Not Available.

Then, we defined the donor age classes. Within each species, the interval between the year of birth of the oldest donor and 2020 was calculated and divided into three equal periods that defined the following classes: VOM (Very Old Material), for donors born in the first third of the interval; OM (Old Material), for those born in the second third; and CM (Contemporary Material) for the recent donors, born in the last third. When possible, we also calculated the donors’ age at their first collection and defined three age classes, SBC, MBC, and EBC, as described in Table 1. SBC ranges from two years after birth (for ruminants and pigs) to five years after birth (for equids), MBC varies from two to four years (for ruminants and pigs) to five to ten years (for equids), and EBC ranges from four years (for ruminants and pigs) to ten years (for equids). Table 1 summarizes the set of indicators used to characterize the FNC collections. Altogether, 12 variables were defined for each donor and six were used to build a classification. The distributions of these values for the six species are available in Additional file 3: Table S2 and Additional file 4: Figure S2. Two collections from different owners were available for the French Landrace pig breed in the FNC and these were analyzed separately.

**Table 1:** Descriptors of the French National Cryobank (FNC) collections.

| Descriptor | Description |
| --- | --- |
| Species* | One of the six mammal species (Donkey, Horse, Cattle, Sheep, Goat, and Pig) |
| Breed | Breed or lines with at least one donor in the FNC |
| Donor | A sire having cryopreserved semen in the FNC |
| Dose | Number of straws of frozen semen required for one artificial insemination according to species and breed |
| Birth year of donors | Year of birth of a donor |
| Year of 1<sup>st</sup> collection | Year of first collection of donor semen |
| Donor's Age at 1<sup>st</sup> collection | Age of the donor at first semen collection |
| Donor's Birth year Class<sup>1*</sup> | Birth year of the donor with semen in collection<br>Very Old Material (VOM)<br>Old Material (OM)<br>Contemporary Material (CM)<br>The values are calculated according to species. |
| Donor's 1st Collection Age Class<sup>1*</sup> | Age group of the donor at the time of its first semen collection<br>Start of the donor's Breeding Career (SBC)<br>Middle of the donor's Breeding Career (MBC)<br>End of the donor's Breeding Career (EBC) |
| Motive (Type) for entry into collection<sup>*</sup> | Reason for entry of a donor into collection, according to the 3 types defined at the creation of the FNC<br>Type I: Donor of an endangered breed<br>Type II: Donor with exceptional characteristics (origins, performance, anomalies)<br>Type III: Representative donor of a breed over a certain period |
| Diffusion<sup>*</sup> | Diffusion status of the breed<br>Local: present only in a specific region<br>National: bred throughout the whole French territory<br>International: with international exchanges of semen or breeding stock |
| Use<sup>*</sup> | Primary use of the breed<br>Meat: raised for meat production<br>Dairy: raised for milk production<br>Wool: raised for wool production<br>Draft_horse: heavy horse breeds originally bred for work and meat<br>Riding_horse: light horse breeds originally bred for sport and leisure activities |

<sup>1</sup> The definition of intervals delimiting the different classes for each species are available in Additional\_file4\_Figure\_S2
\* The variables with an asterisk were used to build regression trees

### Genealogical data on a subset of breeds

We used pedigree data from 17 breeds that represented the diversity of breed management systems encountered in France and that also had several donors in the FNC. The 17 breeds included five pig breeds—Pietrain (PIE), Cul Noir du Limousin (CNL), Duroc (DR), Large White dam line (LWF), and French Landrace (LF)—and five sheep breeds—Dairy Lacaune (LAC), Manech Tête Noire (MTN), Manech Tête Rousse (MTR), Basco-Béarnaise (BAS), and Mouton Vendéen (VEN)—along with seven cattle breeds: Holstein (HOL), Montbéliarde (MON), Abondance (ABO), Froment du Léon (FRO), Tarentaise (TAR), Saosnoise (SAO), and Maraîchine (MAR). Pedigree data from these breeds were used in genealogical analyses, described in detail below.

### Regression trees

We built regression trees to identify the criteria that best explained the data structure in terms of number of donors per breed (Nb_donors_) and number of doses per donor (Nb_doses_). The trees were created and visualized using the rpart package v4.1-15 (Therneau & Atkinson, 2019) and the partykit package v1.2-15 (Hothorn & Zeileis, 2015). They were built using a CART (classification and regression tree) algorithm (Breiman et al., 1984) via recursive divisions of the dataset that aimed to reduce the heterogeneity of the dependent variable at each node. Thus, the partitioning of the dataset was based on minimizing the within-branch variance while maximizing the variance between branches.

In the first model, the number of donors per breed was the dependent variable, while the species, the use (Use), and the diffusion (Diff) of the breed were the explanatory variables.

The first model was described as follows:

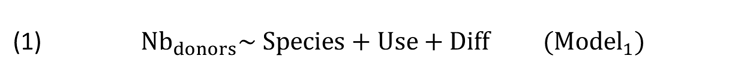

Then, a second model was used to identify the factors that determined the number of doses as a function of species, motivation for entry (Type), breed use (Use), breed diffusion (Diff), and the donor’s birth year class (VOM, OM, or CM) and age class at the first collection (SBC, MBC, or EBC).

It was defined as follows:

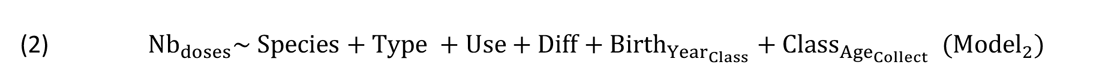

The number of doses and of donors were counts (i.e., without zeros) and thus were assumed to follow a Poisson distribution. The maximum depth of nodes in the final tree was set to three. For each terminal node of the different models, we computed the mean, standard deviation, and percentage of the dependent variable (i.e., number of donors or number of doses).

All analyses were performed using R v4.0.2 (R Core team, 2020) and the graphical representations were created with the ggplot2 package v3.3.5 (Wickham, 2016). Statistical tests were performed with the lm function, post-hoc comparisons were carried out using the emmeans package v1.7.2 (Lenth et al., 2021), type II ANOVAs were performed with the car package v3.0-12 (Fox & Weisberg, 2019), and all correlation coefficients were computed as Pearson correlations with the stats package v4.1.3.

### Assessing the diversity of collections according to species or breed

We adapted the Gini-Simpson index (Gini, 1912; Simpson, 1949) with the goal of measuring the alpha diversity present in the FNC for each of the six species on the basis of the distribution of donors. This index (named E) expresses the probability that two donors from the same species belong to different breeds, and thus enables assessment of the homogeneity of donor sampling across breeds for a given species. It was calculated as follows:

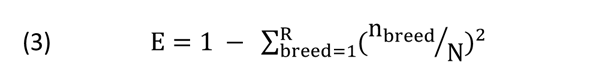

with *n_breed_* the number of donors in the breed, R the total number of breeds in collection for the species, and N the total number of donors of the species.

The optimal value, representing an even distribution of breeds, was calculated for each species according to the following formula:

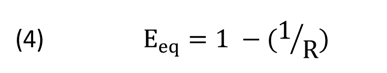

The evenness index Ev, defined as the ratio of E over E_eq_, was calculated to assess how far the collection of a species is from a balanced representation of all breeds.

Then we calculated the effective number of donors per breed, denoted De. This represents the equivalent number of donors in a breed if all doses were distributed equally among donors, and was computed as follows:

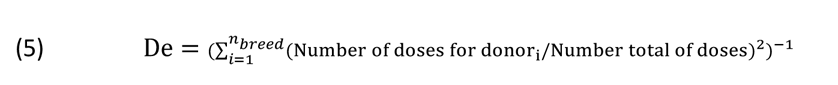

Finally, we calculated the ratio between the effective number of donors and the total number of donors for the breed, to evaluate the unevenness of the representation of donors in a given collection.

### Genetic contribution

For the subset of 17 breeds described above, pedigrees of individuals—either from the breeding scheme for the breed, or from the entire population for breeds in a conservation program—were extracted from herd books or species-specific French national databases. Then, pedigree data were analyzed using PEDIG software (Boichard, 2002). All pedigrees were formatted using the *ped_util* module tracking 25 generations back. The studied populations included the individuals born between 2011 and 2020 and all donors registered in the FNC. Genetic contributions of the FNC donors were calculated from pedigree data using the *contribution* module of PEDIG for each year from 2011 to 2020. The number of direct (i.e., first generation) offspring of the cryoconserved sires was calculated for each year over the same period. We also calculated the annual contribution of the whole set of donors for a breed, along with the individual contribution of each donor.

### Index of Diversity Impact (IDI)

Using the same subset of 17 breeds, we defined contemporary cohorts for each breed that took into account the generation interval for each species, i.e., 3 years for swine (2018-2020), 4 years for sheep (2017-2020), and 6 years for cattle (2015-2020). For females, the cohorts included individuals with at least one known parent. For males, the cohorts included only those males having at least one known parent and one offspring (i.e., sires). The cryobank cohort included all sires with frozen semen for each breed in the FNC. Then, we calculated the average kinship between the different cohorts using the *colleau* module of PEDIG which is based on the inverse of the relationship matrix between individuals as described in Colleau (2022). The kinship coefficient (t) represents the probability an allele drawn randomly from an individual, i, and an allele drawn at the same autosomal locus from another individual, j, are identical and inherited from a common ancestry, i.e. identical by descent (Malécot, 1948). The pedigree quality of these contemporary cohorts was assessed with the *ngen* module of PEDIG using an overall equivalent generation number (EqG), calculated as the weighted average of the equivalent generations for male and female pathways for each year. For each breed with pedigree information, we computed the average kinship within donors in the cryobank, denoted 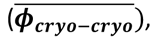 the average kinship within the contemporary male cohort, denoted 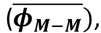 and the average kinship between the cryobank cohort and the contemporary male cohort, denoted 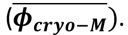 We also computed the average kinship between the cryobank cohort and the contemporary female cohort, denoted 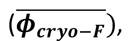 and the average kinship between the contemporary male and female cohorts, denoted 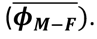 Then, we defined an Index of Diversity Impact (IDI) for the cryobank cohort. This indicator estimates the expected impact on the level of genetic diversity in the next generation if the cryoconserved sires (and only these sires) were used in random mating with contemporary females. The IDI was calculated as follows:

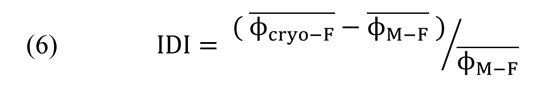

If the males in the cryobank are fully representative of the contemporary population in terms of pedigree relatedness, then IDI is expected to be null. In order to check for deviation from the null value, we calculated 95% confidence intervals (CI) of the IDI as:

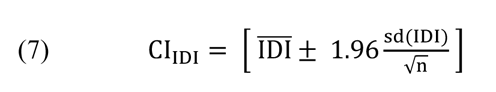

with sd(IDI) the individual standard deviation observed within the cryobank cohort and n the number of males in this cohort.

## Results

### Composition of the French National Cryobank’s collections

#### General findings

The six species represented a total of 115 breeds and 2888 donors with cryopreserved semen in the FNC. The collection includes 21 cattle breeds, 2 donkey breeds, 10 goat breeds, 19 horse breeds, 15 pig breeds or lines, and 48 sheep breeds, respectively corresponding to 977 bulls, 10 jacks, 98 bucks, 175 stallions, 451 boars, and 1,177 rams with frozen semen (Figure 1a).

**Figure 1:**
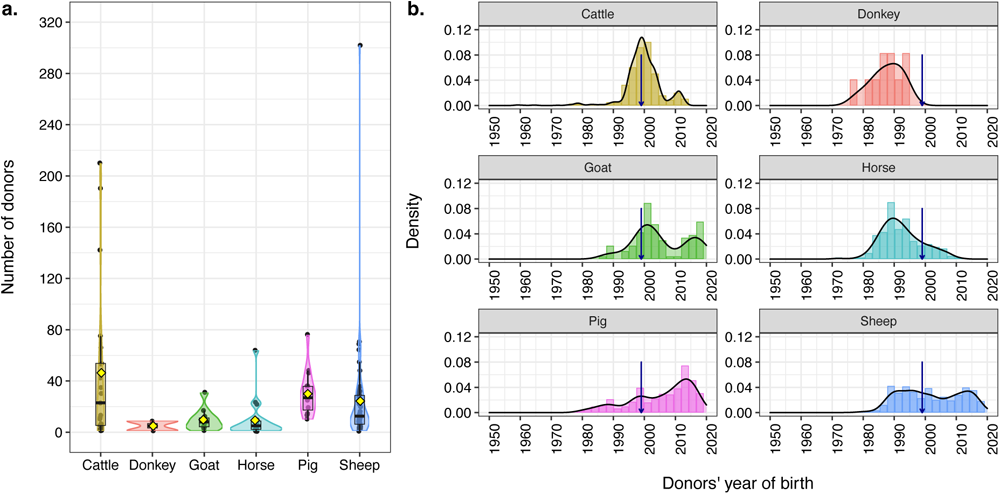
Number of donors per breed (a) and year of birth of donors (b) present in the French National Cryobank for six farm mammal species. The blue arrow represents the creation of the French National Cryobank in 1999.

Donor birth years ranged from 1959 to 2013 for cattle, 1978 to 1994 for donkeys, 1987 to 2019 for goats, 1972 to 2011 for horses, 1979 to 2018 for pigs, and 1974 to 2019 for sheep (Figure 1b). For horses, very few donors were born after the 2000s. In cattle, there were few donors born after 2010. For sheep and goats, donors have been born relatively continuously since 1990. For pigs, there were more donors born between 2010 and 2018. For all species, the collections included material from donors born before the establishment of the FNC, indicating that material from old sires had been retrieved by the FNC after 1999.

Of all donors with frozen semen in the collection, 67.8% belonged to local breeds. Local donkey, horse, and goat breeds represented 17.4% of the donors. Dairy cattle breeds represented 25.7% of the total number of donors, while beef breeds represented 8.1%. Cattle semen represented 52.1% of the total number of doses, compared to 36.2% for sheep, 4.6% for pig, 3.7% for horse, 3.2% for goat, and 0.2% for donkey.

#### Factors determining the number of donors per breed

Node 1 of the regression tree was determined by the diffusion of the breed, with local breeds being separated from the national and international breeds (Figure 2). Within the local breeds branch, node 2 was determined by the species identity, with cattle, pig, and sheep separated from donkey, horse, and goat; this latter group had an average of 8 donors per breed (min=1, max=31). Node 4 differentiated donors from local breeds of cattle, pigs, and sheep according to their use, separating dairy from other uses (meat or wool). Local dairy sheep and cow breeds had an average of 30 donors (min=1, max=75), whereas other local cattle, sheep, or pig breeds had an average of 14 donors (min=1, max=69). For the national and international breeds (node 7 in Figure 2), the first division isolated the dairy breeds; these represented only 6.1% of the breeds in collection, but had a large number of donors on average. Indeed, this branch included the major dairy breeds of ruminants—Holstein, Montbéliarde, Normande, and Brown Swiss for cattle, dairy Lacaune for sheep, and Saanen and Alpine for goats—which had an average of 129 donors each (min=6, max=302). The non-dairy breeds with international and national distribution were then separated by species identity (node 8 in Figure 2).

**Figure 2:**
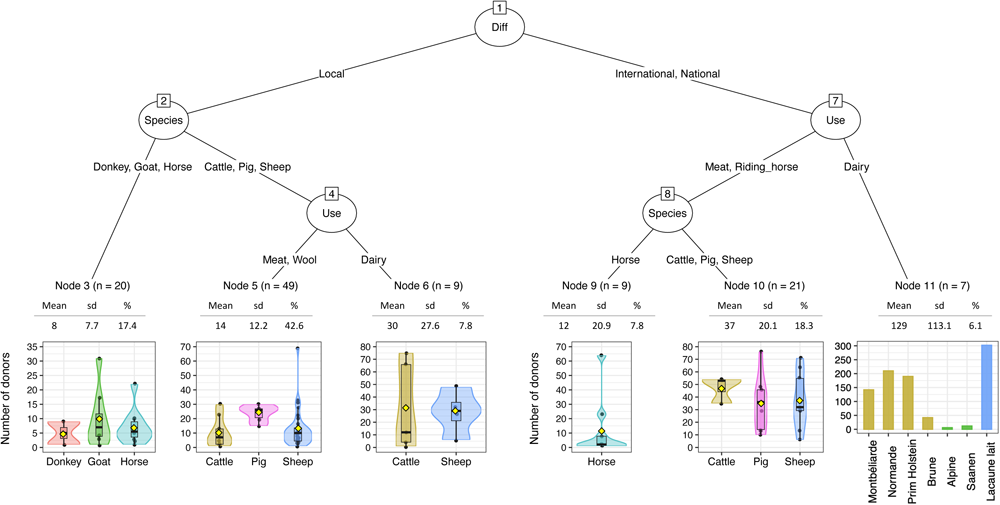
Regression tree for the number of donors present per breed in the French National Cryobank. Diff represents the diffusion of the breed. The percentages represent the proportion of donors present in each terminal node.

The horse breeds had an average of 12 donors (min=1, max=64) whereas the meat breeds from the three other species had an average of 37 donors per breed (min=6, max=76).

#### Factors determining the number of doses per donor

In the regression tree for the number of doses per donor (Figure 3), node 1 separated cattle from the other species. Within cattle, node 7 was defined by the main use of the breed (dairy or meat). In the dairy breed branch, data were further separated (node 8) according to the motivation for entry: types I and II were grouped in node 10, with an average of 199 doses per donor (min=25, max=800), whereas node 9 included only type III, with an average of 104 doses (min=47, max=300). For beef breeds, segregation was based on the diffusion status of the breed (node 11 in Figure 3), separating national breeds (average of 199 doses per donor, min=100, max=300) from local breeds (average of 361 doses per donor, min=183, max=906, se=21.7). Then, in the branch containing the other species (node 2 in Figure 3), pigs were separated from equids and small ruminants, with a lower number of doses for boars (35 on average, min=3, max=92). The other species formed node 4, which was subsequently divided according to the donor’s birth year class. Recent individuals (i.e., CM) clustered in node 6, with an average of 122 doses per donor (min=2, max = 276), whereas older donors (i.e., VOM and OM) clustered in node 5, with an average of 92 doses per donor (min=1, max = 394).

**Figure 3:**
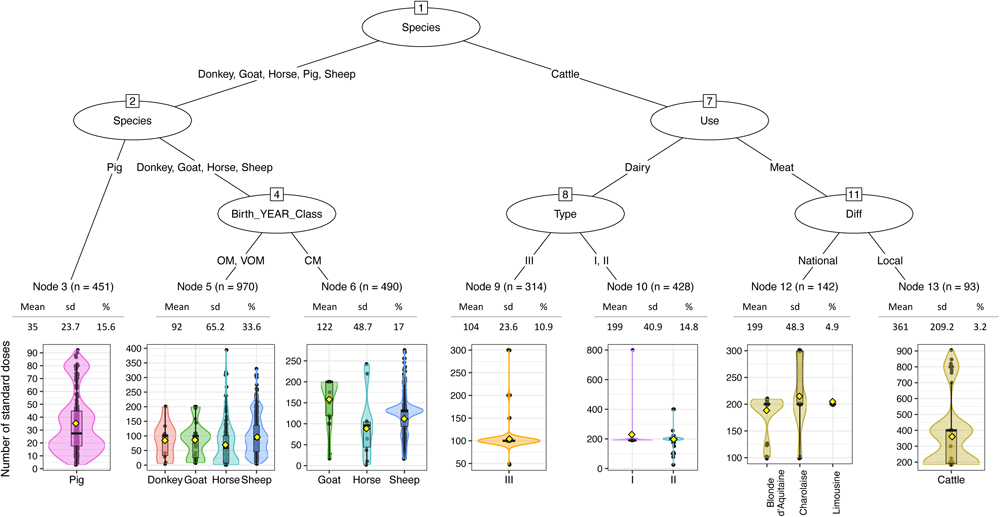
Regression tree for the number of doses per donor present in the French National Cryobank. Birth_YEAR_Class represents the donor’s year of birth as one of three values: VOM (Very Old Material), OM (Old Material), and CM (Contemporary Material).

#### Donor representativeness within species

Gini-Simpson index values varied from 0.18 for donkey to 0.91 for pig and sheep (Table 2). Values of evenness (E divided by E_eq_) ranged from 36.0% for donkey to 97.8% for pig.

**Table 2:** Diversity indicators according to species.

| *Collections* | *Cattle* | *Sheep* | *Goat* | *Pig* | *Horse* | *Donkey* |
| --- | --- | --- | --- | --- | --- | --- |
| *Number of official breeds or lines* | 54 | 59 | 14 | 39 | 54 | 8 |
| *Number of breed/lines in cryobank (R)* | 21 | 48 | 10 | 15 | 19 | 2 |
| Total number of donors | 977 | 1177 | 98 | 451 | 175 | 10 |
| Gini-Simpson index (E) | 0.87 | 0.91 | 0.83 | 0.91 | 0.82 | 0.18 |
| $E_{eq}$ | 0.95 | 0.98 | 0.90 | 0.93 | 0.95 | 0.5 |
| *Evenness index, $Ev=E/E_{eq}$* | 91.7 % | 93.1 % | 91.8 % | 97.8 % | 86.5 % | 36.0 % |

Note: For a fixed number of breeds, $E_{eq}$ corresponds to the value of E if each breed contributed equally

The number of effective donors (De) in the different species ranged from 1 to 248 (Figure 4a). The ratio of the number of effective donors to the number of real donors ranged from 42% to 100% (Figure 4b). Per species, the average ratio between De and actual donor number ranged from 71.6% for horses to 92.0% for cattle. Details on the distribution of donors and doses within the different species and collections are available in Additional file 5: Table S3 and Additional file 6: Table S4.

**Figure 4:**
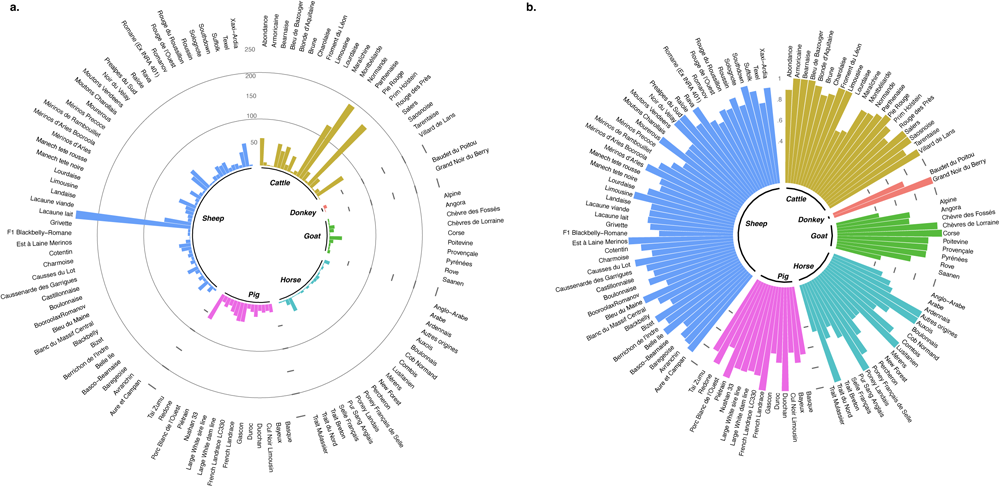
Number of effective donors (De) in the collection (**a**) and ratio between De and the real number of donors in the collection (**b**) in the French National Cryobank according to species. Diff corresponds to the diffusion of the breed. Type is the dominant motivation for cryobank entry: I for endangered breeds, II for individuals with exceptional characteristics, and III for representative individuals of a breed over a given period. The percentages indicate the proportion of donors represented in each terminal node.

### Level of diversity in the collections and comparison with contemporary populations

#### Genetic contribution of germplasm collections in 2020 and evolution since 2011

The genetic contributions of collections to contemporary populations (i.e., 2020) varied between 2.21% for HOL and 63.73% for FRO (Figure 5). We present here data on the evolution over time of those genetic contributions, as well as the number of direct offspring born between 2011 and 2020, for a subset of five breeds differing in selection history and with pedigrees available: HOL and FRO for cattle, CNL for pig, and MTN and VEN for sheep.

**Figure 5:**
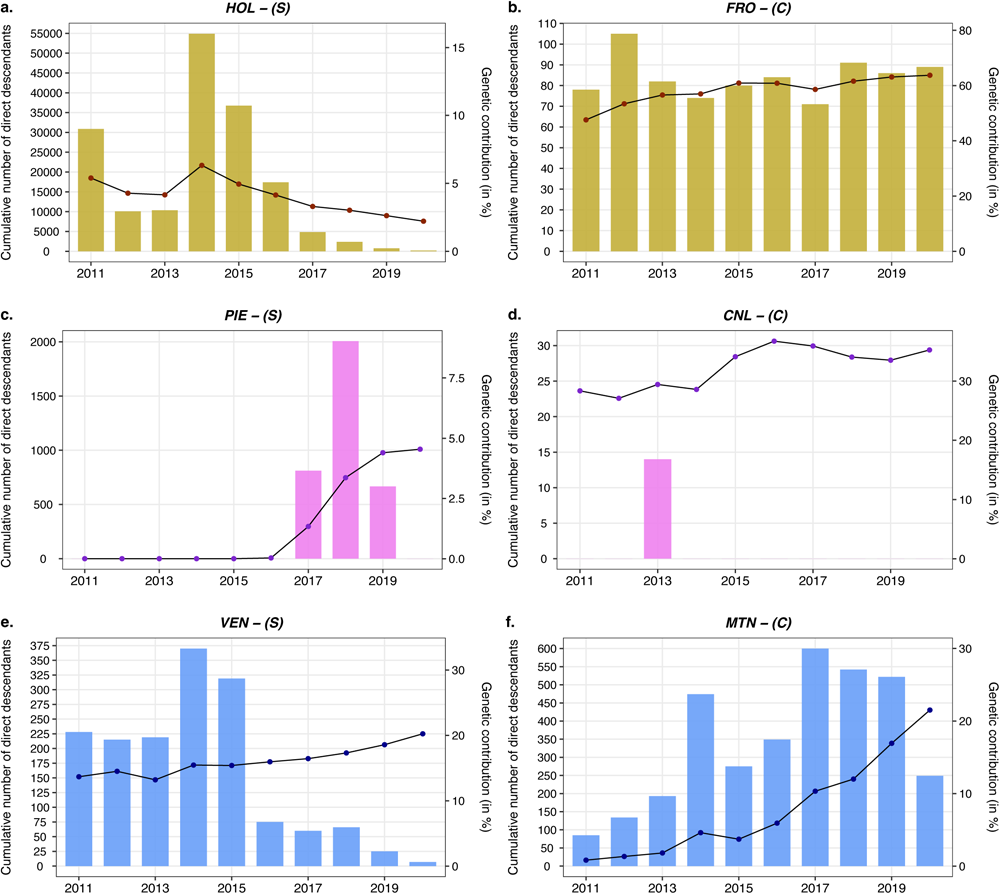
Genetic contribution of the whole collection of cryopreserved donors for different species and breeds differing in their management strategies. The histograms represent the cumulative number of direct offspring and the curves represent the genetic contribution of cryopreserved individuals (in percentage). The abbreviations of the breeds are as described in the Materials and Methods section in the text. The two breeds of dairy cattle are represented in ochre (**a, b**), the two breeds of pigs are represented in pink (**c, d**), and the two breeds of sheep are represented in blue (**e, f**). Large breeds under selection are represented by the letter S and are on the left (**a, c, e**), while breeds with a smaller population size are represented by the letter C and are on the right (**b, d, f**).

Results for the other breeds are shown in Additional file 7: Figure S3. In HOL, the genetic contribution of the collection to the current population has been steadily decreasing since 2014, together with the number of direct offspring from cryobank sires. Instead, in the FRO cattle breed, for which a conservation program is in place, the number of direct offspring from cryobank sires has been relatively stable over the last ten years; all cryobank sires were used, and thus the genetic contribution slowly increased. For the CNL pig breed, although semen from cryopreserved boars in the FNC has not been used recently (except for a few litters in 2013), there has been a slight increase in genetic contribution since 2011. For the two sheep breeds, the genetic contribution of the collections has been steadily increasing since 2014. For VEN, the number of direct offspring has drastically decreased since 2016 due to the age of the rams in the collection (the youngest ram was born in 2012). For MTN, the collections were much more recent, with rams born between 2007 and 2016.

#### Genetic diversity and IDI

The quality of the pedigrees ranged from 3.1 to 22.3 EqG on either the female or male pathway (Table 3). Much deeper pedigree knowledge was available for pigs, which, together with a higher litter size, explains the higher kinship values compared to those observed in the other species. The mean within-cryobank kinship values (Cryo-Cryo) ranged from 0.8% to 24.8%. The average kinship values between cryobank sires and contemporary active males (Cryo-M) ranged from 3.2% to 24.6%, and the mean kinship values within contemporary active males (M-M) ranged from 2.4% to 35.4%. The average kinship values between cryobank sires and contemporary females (Cryo-F) ranged from 3.1% to 24.8%, and the average kinship values between contemporary active males and contemporary females (M-F) ranged from 2.2% to 35.4%.

**Table 3:** Measures of genetic diversity for the 17 breeds with pedigree data.

| Breed | Cryobank sires |  | Contemporary cohort size<sup>1</sup> |  | No. of EqG <sup>2</sup> |  | Average Kinship (in %)<sup>3</sup> |  |  |  |  | Index of Diversity Impact (in %)<sup>4</sup> |  |  |
| --- | --- | --- | --- | --- | --- | --- | --- | --- | --- | --- | --- | --- | --- | --- |
|  | No. | Birth year range | M_active | F | M | F | Cryo-Cryo | M-M | Cryo-M | Cryo-F | M-F | IDI | CI<sub>IDI</sub> | No. donor with negative IDI |
| SAO | 10 | 1991 ;2011 | 70 | 3089 | 3.2 | 3.3 | 0.8 | 2.4 | 3.2 | 3.1 | 2.2 | 43.6 | [8.2;79.0] | 2 |
| FRO | 12 | 1978 ;2006 | 12 | 534 | 3.1 | 3.1 | 10.1 | 11.0 | 11.8 | 12.5 | 11.0 | 13.1 | [-8.4;34.6] | 3 |
| TAR | 75 | 1994 ;2006 | 202 | 26030 | 6.5 | 6.6 | 10.6 | 10.8 | 10.1 | 9.3 | 9.7 | - 4.6 | [-9.4;0.2] | 44 |
| ABO | 66 | 1995 ;2006 | 470 | 71698 | 6.7 | 6.8 | 15.1 | 12.7 | 13.4 | 12.9 | 12.0 | 7.2 | [1.0;13.4] | 26 |
| MON | 142 | 1994 ;2013 | 3658 | 1201388 | 9.4 | 9.5 | 12.1 | 10.8 | 10.8 | 10.8 | 10.8 | 0.0 | [-3.0;3.0] | 67 |
| MAR | 13 | 1989 ;2004 | 67 | 3368 | 6.7 | 6.5 | 5.7 | 8.8 | 8.1 | 7.6 | 8.2 | - 7.8 | [-30.9;15.3] | 8 |
| HOL | 190 | 1993 ;2013 | 10776 | 5069731 | 10.4 | 10.0 | 10.8 | 11.5 | 9.1 | 8.9 | 10.1 | - 12.6 | [-14.7;-10.6] | 153 |
| PIE | 14 | 2016 ;2017 | 401 | 48503 | 19.3 | 19.1 | 12.9 | 12.9 | 12.7 | 12.5 | 12.6 | - 0.6 | [-2.3;1.1] | 9 |
| CNL | 27 | 1981 ;1999 | 14 | 1367 | 15.5 | 15.5 | 24.8 | 35.4 | 24.6 | 24.8 | 35.4 | - 30.1 | [-33.9;-26.3] | 27 |
| DR | 36 | 2006 ;2018 | 179 | 24738 | 15.6 | 15.6 | 8.4 | 7.5 | 3.2 | 2.9 | 7.9 | - 62.9 | [-69.6;-56.2] | 36 |
| LF | 46 | 2001 ;2013 | 526 | 140093 | 22.3 | 22.3 | 14.2 | 18.8 | 15.0 | 15.0 | 18.8 | - 20.2 | [-24.2;-16.2] | 44 |
| LWF | 48 | 2004 ;2015 | 825 | 353941 | 21.7 | 21.7 | 12.6 | 15.6 | 13.7 | 13.7 | 15.6 | - 12.0 | [-16.4;-7.7] | 35 |
| MTN | 26 | 2007 ;2016 | 108 | 11977 | 5.9 | 4.5 | 11.3 | 9.5 | 11.1 | 5.9 | 5.2 | 14.2 | [6.3;22.1] | 6 |
| MTR | 48 | 2008 ;2016 | 616 | 104184 | 9.7 | 8.2 | 6.8 | 7.2 | 7.3 | 5.5 | 5.5 | 0.5 | [-7.5;8.5] | 19 |
| LAC | 302 | 1980 ;2018 | 881 | 708459 | 13.3 | 10.8 | 5.7 | 5.0 | 4.8 | 3.5 | 3.6 | - 3.9 | [-6.7;-1.1] | 115 |
| BAS | 32 | 2009 ;2016 | 261 | 32547 | 8.4 | 6.1 | 10.1 | 11.0 | 10.8 | 6.8 | 7.0 | - 1.9 | [-9.6;5.8] | 12 |
| VEN | 69 | 1989 ;2012 | 306 | 33855 | 13.7 | 10.3 | 5.1 | 6.3 | 5.1 | 3.7 | 4.4 | - 16.3 | [-20.7;-11.9] | 54 |

<sup>1</sup> Contemporary cohorts were defined using the generation interval for each species, i.e., 2 years for swine (2018-2020), 3 years for sheep (2017-2020), and 5 years for cattle (2015-2020), and includes all individuals with at least one known parent in this period.
The contemporary cohorts of females are denoted F. The contemporary cohorts M\_active consider only male individuals having at least one offspring.
<sup>2</sup> Number of equivalent generations computed for the contemporary cohorts on the male (M) and female (F) pathways.
<sup>3</sup> Cryo-Cryo is the average kinship within donors in the cryobank, M-M is the average kinship within the contemporary male cohort, Cryo-M is the average kinship between donors in the cryobank and the contemporary male cohort, Cryo-F is the average kinship between donors in the cryobank and the contemporary female cohort, M-F is the average kinship between the contemporary male cohort and the contemporary female cohort.
<sup>4</sup> IDI is the Index of Diversity Impact and CI<sub>IDI</sub> is the 95% confidence interval.
The abbreviations of the breeds are as described in the Materials and Methods section in the text.

The breeds’ IDI values varied between -62.9% for DR and 43.6% for SAO.

Overall, seven breeds from the three species considered exhibited a significantly negative IDI. This included four pig breeds: the local breed CNL, which had very old boars in collection, and three breeds under selection whose collections include cryopreserved boars born after 2000. The PIE breed, which is under selection, had an IDI close to zero with very recent boars in collection. The three ruminant breeds with a negative IDI were all breeds under selection, including a highly selected bovine breed (HOL) for which the FNC has encouraged the collection of exceptional donors (Type II) as well as old bulls; the sheep breed VEN, for which the donors are very old rams, most born before 2010; and the sheep breed LAC, for which the IDI value was negative but quite close to zero due to a sampling strategy that aimed to represent the genetic variability of the breed. The other ten breeds had a positive or null IDI.

#### Added value of IDI compared to other indicators

Two indicators of the genetic originality of collections are the kinship between cryopreserved males and active males (Cryo-M) or between males within the collection (Cryo-Cryo). Correlations between IDI and each of these indicators were found to be non-significant (respectively, -0.13, t=-0.52, df=15, p=0.61 and -0.35, t=-1.46, df=15, p=0.16).

Among all 17 breeds studied, the correlation between the individual IDI values of donors and their year of birth was significantly positive (0.33, t=12.06, df=1154, p<0.05), but was heavily dependent on the breed (F=68.75, df=1138, p<0.05), and more specifically on the structure of the FNC collection. The distribution of individual IDIs and correlations with birth years are available for each breed in Additional file 8: Figure S4 and Additional file 9: Figure S5.

We also identified a relationship between IDI and the sampling strategy used for collection, i.e., the different motivations for entry, especially for types I and III (F=9.32, df =754, p<0.05). We plotted predicted IDI values for Type I and Type III resources over the last 40 years (available in Additional file 10: Figure S6). Type III donors, which were collected to maintain a picture of the genetic diversity of the population at a given time, had an IDI close to 0, while Type I donors, which belong to endangered breeds, had extreme IDI values, either positive or negative depending on their year of birth.

## Discussion

### Key findings

Here, we propose a standardized framework (Table 4) for the evaluation of cryopreserved collections to assess the diversity and representativeness of collections at different levels. Such an approach will make it possible to compare collections for better management at a larger scale. This framework is based on the use of three indices: i) a version of the Gini-Simpson index adapted for collections (E), which measures the distribution of donors of a species among its different breeds, together with an evenness index (Ev), which represents the extent of deviation from an even distribution of donors across a given number of breeds; ii) the number of effective donors (De), which represents the distribution of doses among donors within a breed; and iii) the index of diversity impact (IDI), which assesses the effect of the use of germplasm collections on the genetic variability of current populations. Compared to other currently existing indicators, IDI provides additional information regarding the genetic contribution of cryopreserved individuals and should be taken into consideration before using frozen semen from the FNC. Furthermore, IDI is unique in that it is oriented toward the use of collections and not only their conservation.

**Table 4:** A possible framework to assess the potential uses of the French National Cryobank.

| *Study level of germplasm collection* | Indicators | Main objectives |
| --- | --- | --- |
| *Materials available in the collection* | Species and breed in collection<br>Type of resources (semen, oocytes, embryos...) | Improve management of collections |
| *Balanced representation of breeds within a species* | Number of donors (D) per breed<br>Gini-Simpson index (E) and evenness index (Ev)<sup>1</sup> | Reconstitute a breed<br>Improve management of collections |
| *Balanced representation of donors within a breed* | Number of doses per donor<br>Effective number of donors (De) and De/D | Reconstitute a breed |
| *Genetic link with populations in the field* | Genetic contribution of collections to the current population<br>Index of diversity impact (IDI) | Improve management of collections<br>Management of *in situ* population diversity |

In order to apply this framework to an analysis of the FNC, we first described the main characteristics and underlying motivations of its collections. The number of donors in collections of the FNC mainly depended on the diffusion of breeds. Local breeds had, on average, fewer donors than breeds with national or international diffusion. Instead, the number of doses per donor depended mainly on a species’ reproductive biology, the use of reproductive technologies, and the breeding context (i.e., selection vs conservation). Among different species and/or breeds, the use of assisted reproductive technologies, in particular artificial insemination with frozen semen, varies greatly in frequency, largely as a result of technical limitations and organizational constraints. The average number of doses per donor was higher and more balanced across donors (De) in cattle than in other species, reflecting the fact that frozen semen has routinely been used for many years in breeding schemes for this species. Similarly, the potential of each collection to contribute to the management of genetic diversity in current populations (IDI) varied widely depending on the status of a breed (i.e., under conservation, selection), its population size, and characteristics of the collection (i.e., date, size, type of material).

### The FNC landscape

For the six species of domestic farm animals studied in this paper, there was a strong heterogeneity in the number of breeds represented per species as well as the number of donors per breed, indicating that approaches for managing genetic diversity differ considerably among breeds or species. For all of these species, however, we found that the collections included donors who were born long before the FNC was established, indicating that the cryobank was able to recover straws collected and stored in other collections, such as private collections.

In terms of the number of breeds represented in the FNC, sheep had the most, with 48, compared to donkeys which only had 2. These numbers are consistent with the number of breeds recognized in France for each species, with 57 breeds for sheep and 7 for donkeys (Verrier et al., 2015).

In examining donors’ years of birth, we found that there are few recent donors in the equid collections. In horses, this can be explained by the end of the public mission of the National Studs in 2010, which aimed to provide access to public stallions via stations throughout France. The end of this initiative meant that the cost of collecting material from a stallion increased, thus decreasing the number of stallions that could be collected on a fixed budget. Through the work of the CRB-Anim infrastructure project, however, there has been some supplementation with stallions born after 2000 (Magistrini et al., 2014). In donkeys, there are still technical obstacles to the use of frozen semen, leading to subfertile matings. This may explain why only two breeds are represented in the FNC. For cattle, few donors were born after 2010, which corresponds to the transition period from progeny testing to genomic selection. With progeny testing, semen from bulls was collected on a massive scale and cryopreserved until the end of the bulls’ genetic evaluation, which required a large population of offspring with performance data and thus a long time period. Unused semen could then be easily sent to the FNC. Since the onset of genomic selection in the 2010s, this is no longer the case. The best males are identified before puberty, and semen is collected, and immediately marketed, from only a subset of these, which limits the availability of (unused) straws for the FNC. This trend is all the more worrying given that some studies have reported a negative impact of the switch to genomic selection on the level of genetic diversity in some breeds (Doekes et al., 2018a; Doublet et al., 2019; Makanjuola et al., 2020). For small ruminants and pigs in the cryobank, the distribution of donor birth years covers a longer range of dates than for the other species. The increase in the number of donors born around the years 2013–2018 can be explained by funding provided by the CRB-Anim infrastructure project to support additional collections. Thus, for pigs, small ruminants, and horses, the CRB-Anim project had a clear impact on the enrichment of the FNC collections.

Going forward, the Gini-Simpson index E and the associated evenness index Ev can be used to identify the breeds for which the collection of additional donors should be prioritized. Equids and goats should be particularly targeted since these species exhibit the lowest E and Ev values. The effective number of donors, De, can be used to target the individual donors (or their close relatives) from whom it would be beneficial to collect additional doses. Again, horse and goat collections should receive particular attention since they exhibited the lowest ratio of De to the total number of donors.

### The main drivers of the FNC collections

Our results highlighted that the number of doses stored per donor in the FNC mainly depends on the reproductive context of the species. Bovines presented the highest average number of doses per donor, which can be explained, as discussed above, by the fact that frozen semen has been routinely used for many years in this species, especially in the dairy sector (Vishwanath, 2003; Lonergan, 2018). For other species or breeds, the preferred techniques are either fresh semen insemination or natural mating, although advances have been made in recent years (Brinsko, 2006; Faigl et al., 2012; Didion et al., 2013). The main limitations to the use of frozen semen are: i) in pigs, higher cost and decreased fertility (Gonzalez-Peña et al., 2014; Waberski et al., 2019), ii) in donkeys, failed reproduction in jennies (Oliveira et al., 2006) despite successful mule production in mares (Vidament et al., 2009; de Oliveira et al., 2016), and iii) in ovines, the requirement for laparoscopy and abdomen incision for insemination (Souza et al., 1994; Cseh et al., 2012).

Our study revealed that local breeds generally have a lower number of semen donors in the cryobank than national and international breeds. This difference can be explained by both technical considerations and reproductive practices. The use of frozen semen is more frequent in national and international breeds thanks to a set of techniques and procedures that make it easier to collect donors and distribute their semen (DeJarnette et al., 2004; Funk, 2006). In local breeds, instead, frozen semen is more rarely used because sires are distributed over smaller and potentially more remote geographical areas, which tends to make semen collection more costly and difficult. Another obstacle may be requirements related to sanitary regulations, particularly in the French overseas departments and territories, or for endemic breeds in Corsica. For local breeds, the sanitary constraints are associated with high costs and burdensome protocols that slow down the entry of individuals in collection centers. However, there are exceptions to this pattern, since some local breeds do exhibit a high number of donors, sometimes even higher than the national breeds. This is the case for goats, where the two national breeds, Alpine and Saanen, have 6 and 12 donors in the FNC, respectively, while the local breeds Chèvres des Fossés and Poitevine have 17 and 31 donors, respectively. This apparent paradox can be explained by the collection strategy in recent years. Bucks produce much fewer doses than bulls, which means that, in the two national breeds, there are fewer doses left over for preservation. For some threatened local breeds, though, technical and financial resources have been deployed to collect different males for the FNC collections, such as in the CRB-Anim project. Therefore the number of donors in the FNC mainly depends on the economic landscape of the breeding sector of a species.

From the perspective of using the genetic resources present in the FNC, the FAO has published recommendations regarding ideal sperm concentrations and cryopreservation protocols for different species (FAO, 1998), along with guidelines for the numbers of donors and doses needed to reconstitute an extinct population (FAO, 2012). For example, for ruminants and horses, a population recovery based on 150 founder females and 25 cryopreserved males would require the use of 31 to 36 doses per male (for a pregnancy rate of 0.4 to 0.6, respectively). Overall, our work indicates that the FNC collections do not yet meet these recommendations: only 32 of 103 breeds have enough semen doses available for breed reconstitution (10 of 21 cattle breeds, 14 of 48 sheep breeds, 1 of 10 goat breeds, 6 of 15 pig breeds, and 1 of 19 horse breeds, see Additional file 6: Table S4). To comply with FAO recommendations, the FNC has developed targets for the number of doses to conserve for each species. For instance, the target for pigs was to have 25 boars with 40 doses in order to have a better representation of the populations under selection, which explains the higher number of doses included for donors born after 1995. For the very old boars in the collection, semen was collected before the implementation of these recommendations, which explains the lower number of doses available. For both small ruminant and equine species, the number of doses present also depends on the year of birth of the donors. Contemporary donors, i.e., those born after 2005 for small ruminants and 2004 for horses, have a higher number of doses than older donors, which can be explained by the technical progress made in reproductive biotechnologies and a better control of the donor population. However, some studies have reported that there is also individual variability affecting the performance of frozen semen (Quintero-Moreno et al., 2004; Loomis & Graham, 2008; Contri et al., 2012). In all species, the quality of semen for some sires is impaired by the freezing-thawing process, which affects the potential usefulness of collections. For dairy cattle, in which insemination with frozen semen has become routine, the post-thawing quality of the semen is tested and used to eliminate the bulls with unsatisfactory results. For the ovine collection, the number of doses and rams with semen in the cryobank is higher for types I and III, because of increased efforts for the conservation of local breeds (type I) or for a better representation of the genetic variability of the population (type III). Instead, type II samples characterize unique individuals chosen on the basis of their breeding values, their genetic originality, or the presence of specific alleles, an approach that requires access to data that are available only for a few sheep breeds. For sheep, the number of doses available is also linked to the age at collection: the majority of semen collected came from individuals that were between 18 months and 5 years old. Ejaculates from very young or old rams will produce fewer doses due to different quality factors in the semen (Štolc et al., 2009; Benia et al., 2018).

Overall, continuous improvements in semen quality and freezing-thawing protocols are a major objective of the FNC. Indeed, for recently collected donors the quality of semen does comply with the most recent technical requirements, taking into account differences between species.

### Diversity indices as a guide for using the collections

The IDI index provides a quick synthesis of the potential value of a cryopreserved collection for managing the genetic diversity of the current population of a given breed. It is applicable to all species and breeds, provided that genetic relationships can be estimated between cryoconserved sires and the contemporary population. As shown by the comparisons with the different kinship coefficients, the IDI brings additional, easily interpretable information to analyses of cryobank collections. It provides a snapshot of the value of cryopreserved material to a breed, which here highlighted notable differences, especially among local breeds with small population sizes. Positive IDI values, indicating a negative impact on genetic diversity, were obtained when the collection was either recent (as in the MTN breed) or represented most if not all of the sire origins in the breed (as in the FRO breed), so that genetic relationships between cryoconserved sires and contemporary females were high. In this case, using the FNC collection may not add much diversity to the population. However, one of the motivations for safeguarding genetic resources for small populations is their vulnerability to epizootic diseases. Thus, even in the case where IDI values are not favorable for immediate use, the situation could be reversed in a few generations only, in the case of population reduction events. Instead, when the collection is old (as in the local breeds MAR or CNL), a negative IDI was obtained, with some extremely low values revealing a significant “kinship gap” between the cryopreserved males and the current population. In pigs, certain commercial lines within a breed are sometimes managed by different breeding companies (Hulsegge et al., 2019; Bidanel et al., 2020), leading to greater genetic distances between some cryopreserved boars and current populations, and thus negative IDI values. For breeds under strong selection (such as HOL or LWF), negative IDI values are the result of rapid genetic gain and a consequent increase in genetic differentiation between cryopreserved sires and current sires. In such a case, using cryoconserved sires could limit inbreeding in the next generation of the contemporary population but would negatively affect the traits under selection, which is the main limiting factor for the use of cryopreserved material in breeding schemes (Leroy et al., 2011; Jacques et al., 2023). However, recent simulation studies reported that a satisfactory compromise could be obtained for both genetic merit and diversity by optimizing the choice of cryopreserved sires (Eynard et al., 2018; Doekes et al., 2018b) and by using preferential matings of these sires with elite females (Dechow et al., 2020; Jacques et al., 2023). Beyond the mean IDI value of a collection, it will also be useful to look at individual IDI values to see if some donors behave as outliers, either by having a stronger genetic contribution to the current population than others (positive individual IDI despite negative average IDI) or by having a rare origin (negative individual IDI despite positive average IDI). The IDI is calculated from pedigree data, so the quality of pedigrees determine the reliability of index values. However, to ensure for IDI estimation reliability, one should have access to pedigrees deep enough only for contemporary populations in order to retrieve a potential relatedness with cryoconserved population. Reversely, this latter individuals can be described by only few generations, the rest being negligible in particular if contemporary individuals are far from cryopreserved ones in the pedigree. Nevertheless, in breeds with low pedigree depths, the use of molecular data is advisible (see below).

In general, very ancient sires may bring back alleles that have been lost over time, either due to selection or to genetic drift. A detailed molecular characterization of the genomic diversity of cryopreserved individuals could allow us to identify these alleles. Moreover, this could create the possibility of calculating IDI from molecular relationships, making it applicable for the evaluation of FNC collections of breeds that do not have complete pedigree data, in particular for local breeds. This would require genomic data from cryopreserved individuals, as well as a representative sample of the genetic diversity of the contemporary population. In addition, calculations of IDI cannot be simply applied to most kinship or relatedness estimates based on molecular data. Indeed, in most cases (if not all when ignoring population structure), the subsequent coefficients are not probabilities and they can be negative, in particular with classical Genomic Relationship Matrix, GRM (Goudet et al., 2018). This makes both computation and interpretation more tricky. Thus, when pedigrees are not available, IDI computation involves to compute molecular based estimates of kinship coefficients by either, i) using a method estimating the relatedness as a proportion of IBD (e.g. KING software by Manichaikul et al., 2010) and then setting to 0 the negative kinship estimates, or ii) reconstructing pedigrees from molecular data to estimate probability based kinship coeeficients for instance using the sequoia package (Huisman, 2017). In this latter case, one should have access to enough genotyped individuals to retrieve a complete enough pedigree to get reliable kinship estimates which is rarely the case for populations without any pedigree data. Unfortunately, genotypes are still a rare commodity for most species, in particular for females, with the exception of certain cattle breeds. Despite this, recent studies highlighted the various opportunities that genomic approaches could create for the characterization and use of cryopreserved genetic resources in gene banks (Oldenbroek & Windig, 2022; Ajmone-Marsan et al., 2023). Although previous studies have assessed the diversity present in gene bank collections (Blackburn, 2018; van Breukelen et al., 2019; Mercat et al., 2020), the IDI makes it possible to link cryobank diversity with that in current populations. It is therefore an index that highlights the value of collections for diversity management for a variety of reasons, both in the case of local breeds and international breeds. This new index could also be regularly updated for germplasm collections in order to follow their progress and to help guide the management strategy of a gene bank, i.e., to prioritize the collection of certain breeds or sires depending on the genetic trends in the standing populations.

## Conclusions

In conclusion, the FNC contains a great diversity of material for different species and breeds of farm animals, both in terms of the quantity of cryopreserved material and of the level of genetic diversity compared to that present in current populations. The framework developed in our case study enables a thorough evaluation of available resources and the characterization of their potential usefulness for managing genetic diversity in animal breeding programs based on different diversity indices. The new diversity index introduced here, IDI, helps to quickly and comprehensively assess the potential of cryopreserved breed collections to reintroduce genetic diversity in current populations. For breeds under strong selection, collection material could be useful in bringing back diversity that can be lost over generations, while for breeds with small population sizes, like local or endangered breeds, particular attention must be paid to the choice of preserved semen, since ancient sires can be the main genetic contributors to the current population. As discussed by numerous previous studies, it is important to identify strategies for the collection of genetic resources in gene banks, but also for the distribution of material. The IDI index can be used to inform both distribution and collection (i.e., whether new material should be representative of the current population or unique). Currently, there is very little use of the genetic resources present in gene banks due to technical and financial limitations. Furthermore, we think that the lack of recommendations and guidelines for their use don’t encourage the careful use of these collections, which remain depletable resources if collections are not restored. More studies are needed to help population managers identify the utility and the limits of the use of gene bank resources. Moreover, for some species, the use of frozen semen remains a delicate and difficult technical operation, and it is important to have a full picture of the possible gains that might be achieved by using individuals from cryobanks. The individual variability in fertility in thawed semen can strongly limit the expected benefit. Further research in reproductive biotechnology is still needed in species in which frozen semen is not routinely used in order to help mobilize these genetic resources and conserve genetic diversity.

## Acknowledgments

The authors would like to acknowledge all the managers of the studied breeds who sent us useful information for this study. We would like to thank the Centre Départemental de l’Elevage Ovin des Pyrénées-Atlantiques (CDEO), the organization of approved selection of the Lacaune (UPRA Lacaune), Alliance R&D, and LIGERAL, who permitted us to have access to the data and provided comments concerning their breeds. The authors would like to acknowledge Diane Buisson and Jean-Baptiste Tertrain for their help in extracting the pedigree data. The authors also thank the staff of the French National Cryobank and the partners of the CRB-Anim infrastructure project, ANR-11-INBS-0003, funded by the French National Research Agency through the ‘Investing for the Future’ program, for all the information made available for this study.

## Data, scripts, and code availability

The datasets and scripts used in this study are available at https://doi.org/10.5281/zenodo.8163138.

## Supplementary information

Additional files are available below as well as online at https://doi.org/10.5281/zenodo.8162989:

Additional file 1: Table S1

**Format: .xlsx**

**Title:** Report on the output of material since the creation of the French National Cryobank

Additional file 2: Figure S1

**Format: .xlsx**

**Title:** Summary of material outputs from the French National Cryobank since 1999

Additional file 3: Table S2

**Format: .xlsx**

**Title:** Descriptors from the French National Cryobank data for six species

Additional file 4: Figure S2

**Format: .pdf**

**Title:** Distribution of data and definition of intervals regarding breed diffusion (A), donor birth-year classes (B), and classes of donor’s age at 1^st^ collection (C)

Additional file 5: Table S3

**Format: .xlsx**

**Title:** Statistics of the number of effective donors (De) according to species

Additional file 6: Table S4

**Format: .xlsx**

**Title:** Distribution of donors and doses across breeds

**Description:** In green, the breeds for which the FNC collections meet FAO conditions for reconstitution of an extinct breed. In blue, the breeds for which the collections of the FNC do not yet meet FAO conditions for reconstitution of an extinct breed. In gray, FAO recommendations not available. Based on the 2012 FAO report, with 100 founder females for ruminants and horses, or 30 founder females for pigs (with a pregnancy rate of 0.6).

Additional file 7: Figure S3

**Format: .pdf**

**Title:** Evolution of genetic contributions of sires and their production of direct descendants in the French National Cryobank over the period 2011–2020 for three livestock species

**Description:** Two breeds of pigs are represented in pink (a, b), three breeds of sheep are represented in blue (c, d, e), and five breeds of cattle are represented in ochre (f, g, h, i, j).

Additional file 8: Figure S4

**Format: .pdf**

**Title:** Distribution of individual IDIs of cryopreserved sires for the 17 breeds analyzed

**Description:** Pig breeds are represented in pink, sheep breeds are represented in blue, and cattle breeds are represented in ochre.

Additional file 9: Figure S5

**Format: .pdf**

**Title:** Correlation between IDI values and the year of birth of donors

Additional file 10: Figure S6

**Format: .pdf**

**Title:** Prediction of IDI values based on donor’s year of birth and the motivation for entry into collection

**Description:** Type I (for endangered breeds) is represented in purple and type III (for representative individuals of a breed over a period) is represented in orange.

## Authors’ contribution

GR and MTB conceived the project. DD helped with data recovery for the FNC. MJM and CDB helped to obtain pedigree data. AJ performed the analysis and drafted the manuscript. GR and MTB supervised the analyses. AJ, MTB, and GR interpreted the results. AJ, GR, DD, CDB, MJM, and MTB contributed to the writing of the manuscript. All authors read and approved the final manuscript.

